# Genetic consequences of improved river connectivity in brown trout (*Salmo trutta*, L.)

**DOI:** 10.1101/2023.07.30.551162

**Authors:** Paolo Moccetti, Jamie R. Dodd, Domino A. Joyce, Andy D. Nunn, Ben Gillespie, Jonathan D. Bolland

## Abstract

Fragmentation of watercourses poses a significant threat to biodiversity, particularly for migratory fish species. Mitigation measures such as fishways, have been increasingly implemented to restore river connectivity and support fish migration. The effects of such restoration efforts are typically tested using telemetry and fisheries methods, which do not fully capture the broader population movements that may have important consequences for population viability. We performed a before-and-after control-impact (BACI) study using genetic tools (SNPs) to investigate the effect of a newly implemented fishway, aiming to enhance upstream spawning migration of brown trout (*Salmo trutta*, Linnaeus) in a reservoir with two headwater tributaries fragmented by man-made weirs. Another reservoir with two barrier-free tributaries was also analysed as a control. Our results showed that the isolated brown trout population was spawning in the reservoir before the installation of the fishway, and we found genetic structuring and differentiation between fragmented headwater tributaries before the fishway construction, but not in the control reservoir. Unexpectedly, after the fishway construction we observed signals consistent with increased genetic differentiation between populations of newly recruited juvenile fish in the reservoir tributary and fish in the reservoir. We propose this was caused by newly enabled philopatric behaviour of brown trout to their natal spawning tributary. In contrast, we did not find any genetic changes in the tributary without a fishway or in the barrier-free reservoir system. Given the scarcity of similar studies, we advocate for an increased use of genetic analyses in BACI studies to monitor and evaluate the effect of efforts to restore habitat connectivity and inform future management strategies.

## Introduction

Fragmentation and loss of habitat caused by the conversion of land and waterways for human activities is a serious threat to Earth’s biological diversity (Collinge, 1996; Jetz, Wilcove & Dobson, 2007). While fragmentation and restoration in terrestrial habitats have received considerable attention, the impact of watercourse fragmentation has been neglected until recently, despite its profound effects on hydrology, geomorphology, biodiversity, water quality, and the well-being of human communities tied to these water systems (Fuller, Doyle & Strayer, 2015; Jumani et al. 2020). For centuries, river systems have been altered for industry, including power, agriculture and flood control, through the construction of dams and weirs (Benke, 1990; Vitousek et al., 1997; Liermann et al., 2012). This infrastructure has caused a reduction in the connectivity of riverine ecosystems (Cowx and Welcomme, 1998), delayed migration of fish species (Caudill et al., 2007) and in some cases made sections of river completely impassable (Clark et al., 2020). It is estimated that there are at least 1.2 million river-barriers in Europe (Belletti et al., 2020), with almost 66,000 obstructions in Great Britain alone (Jones et al., 2019), that are either restricting or halting access to key life stage habitats for native fish species. These obstructions have a detrimental effect on fish communities (Belliard et al., 2009; Brown et al., 2013; Harris et al., 2017) causing a number of riverine fish species to become critically endangered or extinct (Nicola, Elvira & Almodóvar, 1996; Humphries & Winemiller, 2009; Liermann et al., 2012; Dias et al., 2017; Pavlova et al., 2017). Anthropogenic fragmentation in fish populations can also lead to genetic erosion, increased risk of inbreeding and reduced potential for genetic adaptation (Bijlsma & Loeschcke, 2012; Stelkens et al., 2012; Coleman et al., 2018; Vera-Escalona et al., 2018; González-Ferreras et al., 2022).

In recent decades, in North America and Western Europe there has been a growing commitment to restore riverine connectivity by removing artificial barriers (Habel et al., 2020), often associated with significant financial costs. Fishways, structures designed to enhance fish passage, have also been extensively employed as a solution to remediate the negative influence of fragmentation in cases where the complete removal of physical barriers is not achievable (Dodd, Cowx & Bolland, 2017). However, the effectiveness and impact of fishway structures is highly variable, highlighting the importance of case-specific testing and evaluation (Noonan, Grant & Jackson, 2012). Evaluation of fishway effectiveness in enhancing fish movements and restoring longitudinal connectivity typically involves fish telemetry, video and acoustic camera monitoring and population abundance surveys over time (Bunt, Castro-Santos & Haro, 2012; Birnie-Gauvin et al., 2017; Pennock et al., 2017; Dodd, Cowx & Bolland, 2018; Wilkes et al., 2019). While these methods are valuable in understanding the individual movement patterns of fish in relation to fishway characteristics and changes in recruitment due to improved habitat connectivity, they have technical, spatial, and temporal limitations (Wilkes et al., 2019). They do not fully capture the broader population movements and the origin of fish using the fishway, and therefore the longer-term population changes that might have important consequences for population longevity. For instance, accounting for gene flow between partially isolated populations is crucial, as it can mitigate the decline in genetic diversity and reduce the risk of extinction by connecting these populations.

Incorporating genetic analyses into the assessment of the effects of fishways and barrier removal can provide a more comprehensive understanding of gene flow, genetic diversity, population structure, and the long-term viability and resilience of wild fish populations, as recommended by Wilkes et al. (2019). Most studies, though, have focussed on investigating the contemporary genetic structure either in situations of existing river fragmentation or once measures to restore river connectivity have been implemented (e.g., Meldgaard, Nielsen & Loeschcke, 2003; Kinziger et al., 2008; Perrier et al., 2014; Torterotot et al., 2014; Carim et al., 2016; Gouskov et al., 2016; Ferreira et al., 2017; Van Leeuwen et al., 2018; González-Ferreras et al., 2022), largely due to limited availability of usable tissue samples (for DNA extraction) collected prior to the intervention (Zarri et al., 2022). This makes it difficult to quantify the changes that are a result of the intervention. Before-and-after studies directly comparing genetic changes following barrier removal and river restoration are the best way to test the effect of mitigation measures, but they are almost absent in the literature (Zarri et al., 2022), with just a few studies attempting to investigate the genetic effects of restoration interventions on fish (Neville, Dauwalter & Peacock, 2016; Wood, Welsh & Todd Petty, 2018; Fraik et al., 2021). These studies exclusively focussed on barrier removals, such as culverts and dams, neglecting the investigation of fishways. Fishways are potentially less effective in enhancing fish migration compared to actual removals and may also impose selective pressures on specific fish types (McLaughlin et al., 2013; Silva et al., 2018; Tamario et al., 2019; Wilkes et al., 2019; Zarri et al., 2022). Furthermore, the difficulty in identifying suitable “control” populations for comparison mean no genetic studies are yet to perform before-after control-impact (BACI) analysis, despite being recognised as the most robust approach to test the effect of an impact variable (Moland et al., 2021). Indeed, while comparisons of fragmented and restored watercourses can provide general insights into effects of habitat restoration actions, the BACI approach yields specific and practical details regarding the removal or mitigation of particular structures within a river, and more broadly when assessing the impact of connectivity management measures in conservation. The scarcity of such studies hinders scientists, managers and decision-makers from truly assessing the efficacy of measures to mitigate habitat fragmentation. Our study addresses this using brown trout (*Salmo trutta*, Linneaus) before and after the installation of a fishway in comparison to those in a fragmented tributary and in two barrier-free tributaries as controls.

Brown trout are a globally distributed species known to exhibit high variability in life history and movement strategies, which encompass residency, potamodromy (freshwater migratory) and anadromy (migration between freshwater and sea) (Klemetsen et al., 2003; Klemetsen, 2013; Ferguson & Prodöhl, 2022). These strategies involve spawning, feeding and refuge migrations within their life cycle (Banks, 1969; Jonsson and Jonsson, 2011; Ferguson et al., 2019). Brown trout typically spawn on gravel substrate in running water (Klemetsen et al., 2003) but there are infrequent reports of lake spawning (Brabrand, Koestler & Borgstrøm, 2002; Brabrand, Hansen & Koestler, 2006; Arostegui & Quinn, 2019). Like most salmonid species, brown trout exhibit philopatric behaviour, i.e. normally ‘home’ to their natal river and nursery grounds to spawn (Jonsson & Jonsson, 2011; Keefer & Caudill, 2014). This makes them highly susceptible to the impacts of riverine barriers, especially in populations where a subset of adults undertakes an obligatory upstream migration from lentic or marine locations. The consequences of anthropogenic fragmentation on brown trout population structure have been partly investigated (e.g., Stelkens et al., 2012; Klütsch et al., 2019; González-Ferreras et al., 2022), but the impact of fishway construction and barrier removal remains largely unknown, partly due to the complexity of their life history strategies. Also, the study of non-anadromous salmonid species often receives considerably less attention compared to their anadromous counterparts, possibly due to the higher economic and recreational value of the latter (Silva et al., 2018; Ferguson et al., 2019; García-Vega et al., 2022). This bias has resulted in a lack of tools and knowledge for managing the freshwater migratory forms of brown trout, which are declining in several European countries (Waldner et al., 2020).

Here, we performed one of the first before-after control-impact genetic investigations to evaluate the impact of constructing a fishway on a wild fish population. We collected samples from a (control) reservoir and two tributaries with no artificial barriers, and a second reservoir (impact) and two tributaries which had large barriers (weirs) that prevented brown trout migration from the reservoir (Dodd et al., 2023). It was unknown whether brown trout could spawn and recruit in the reservoir or if the fish in the reservoir originated from downstream movements from the headwater tributaries.

Samples were collected from both reservoir systems in the three years before and up to five years after the construction of a fishway on one of the reservoir tributaries with a large weir. We used a KASP genotyping assay to generate SNP data (81 loci) to examine the genetic consequences of historic fragmentation, i.e. before data, and efforts to restore river connectivity, i.e. after data, on brown trout. Specifically, we asked whether (I) barriers had promoted genetic structuring and low genetic diversity within the impact reservoir system, and (II) enhanced upstream migration of brown trout through the fishway would result in a decrease in genetic structuring and an increase in admixture within the tributary, compared to both the no barrier tributary, and the no barrier control system.

## Material and Methods

### Study system

Langsett (53°29’47.0’N, 1°41’10.3’W) and Grimwith (54°04’38.6’N, 1°54’36.2’W) reservoirs are geographically isolated (∼ 66 km apart) artificial impoundments situated at 245 and 288 m, respectively, above sea level in the north of England, UK (Fig. 1 A). The original impoundments were built in 1904 (Langsett) and 1864 (Grimwith), with the latter being increased in size in 1983. They have surface areas of 0.49 and 1.43 km^2^ and mean depths of 12.65 and 14.8 m, respectively. Both reservoirs have a simple fish community and harbour land-locked brown trout populations. Additionally, Langsett Reservoir harbours three-spined stickleback (*Gasterosteus aculeatus*, Linneaus), while Grimwith Reservoir harbours Eurasian minnow (*Phoxinus phoxinus*, Linneaus) and rainbow trout (*Oncorhynchus mykiss*, Walbaum), stocked for recreational fishing.

**Figure 1.**
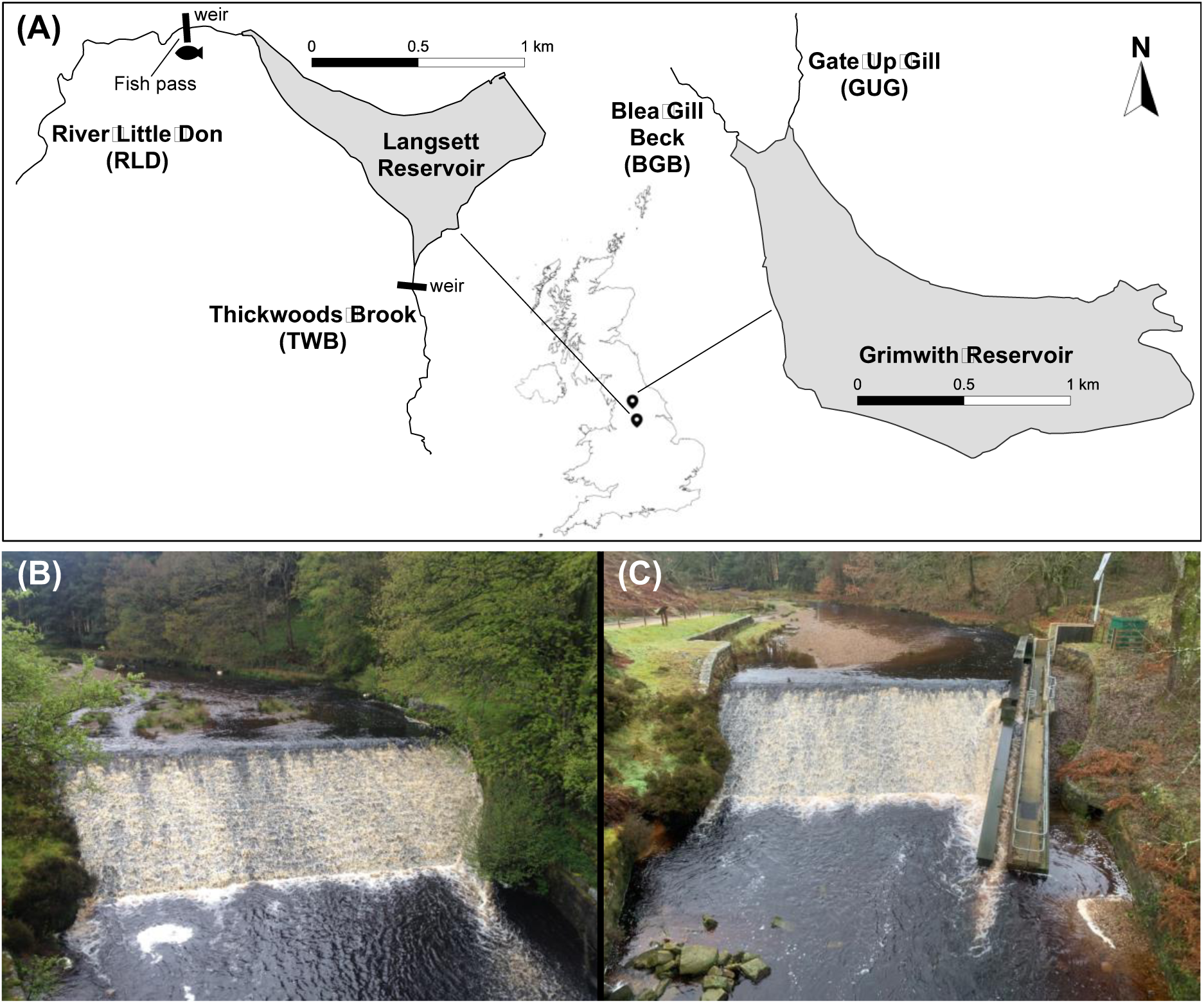
Study area map of Langsett and Grimwith reservoir systems (A) and weir on the River Little Don (RLD) at Langsett, before (B) and after (C) the Larinier fish pass was built in December 2017.

Langsett Reservoir has substantial weirs built in 1904 alongside the impoundment situated on the mouths of the only two viable spawning tributaries, the River Little Don (hereafter RLD) and Thickwoods Brook (hereafter TWB) (Figs. 1 A, B). Tracking studies carried out between 2014-2017 showed that these weirs prevented the upstream migration of brown trout from Langsett Reservoir to the headwater tributaries, but could not definitively determine whether individuals were able to spawn within the reservoir (Dodd et al., 2023). However, there was evidence of fish moving downstream from the headwater tributaries into the reservoir (Dodd et al., 2023). In December 2017, a Larinier-style fish pass (21.7m x 0.45m, 1:7.9) was opened on the RLD weir (Figs. 1 B, C) to ease passage for migrating brown trout from Langsett Reservoir into the headwater tributary, whereas no fishway was installed on the TWB tributary. With the exception of 2017 (when the fish pass opened in December), the fish pass was operational between 1 October - 31 April each year, coinciding with brown trout spawning migration. Tracking analyses from 2017-2021 detected upstream movements of brown trout through the fish pass with overall passage efficiency varying between 7 - 78% (Dodd et al., 2023).

The two tributaries in Grimwith Reservoir, Blea Gill Beck (hereafter BGB) and Gate Up Gill (hereafter GUG), lack barriers preventing migration (Fig. 1 A). Therefore, Grimwith system was used as a baseline comparison for assessing the effects of fragmentation and subsequent restored connectivity on brown trout populations within Langsett system, after construction of the fish pass on the RLD.

### Sampling

Fisheries surveys were performed annually in the main tributaries and reservoirs at Langsett and Grimwith systems in September-October 2015 – 2022 (see Table 1, Suppl. Tables 1-2, and Suppl. Figs. 1, 2, for details about sampling years, sites and fish length distributions). For subsequent analyses, fish sampled between 2015 and 2017 were aggregated and categorised as part of the ‘Before’ dataset, and those sampled between 2018 and 2022 were included in the ‘After’ dataset (Table 1). For the ‘After’ dataset, we included as many young-of-the-year fish as possible in order to include multiple generations and capture genetic changes that were not a result of migration of older fish among locations (Suppl. Figs. 1, 2). Brown trout were captured using a combination of seine and fyke netting in the reservoirs (seine: approximately 150 m x 4 m; 20 mm stretch mesh - set from a boat in a rectangle parallel to the bank and hauled ashore; double fyke nets, 10, 14 and 17 mm mesh size, 53 cm ring size, 2.75 m net length, 6 m ‘leader’ net attached to both ends) and backpack electric fishing equipment (Electracatch 24 V DC input, 200-400 V, 100 W, 50 Hz Pulsed DC, variable pulse width output) in the tributaries. Following each sampling event, captured fish were transferred to water-filled containers where they were monitored, before being measured (fork length, mm) and scale samples were removed and stored in paper envelopes at room temperature. In 2022, a ventral fin-clip was also taken from every fish and stored in absolute ethanol for later DNA extraction. After processing, all fish were then held in a recovery tank until they regained balance and were actively swimming before being returned to the river/reservoir at the site of capture. All fish were treated in compliance with the UK Animals in Scientific Procedures Act 1986 under Home Office licence numbers PPL 60/4400 and PD6C17B56.

**Figure 2.**
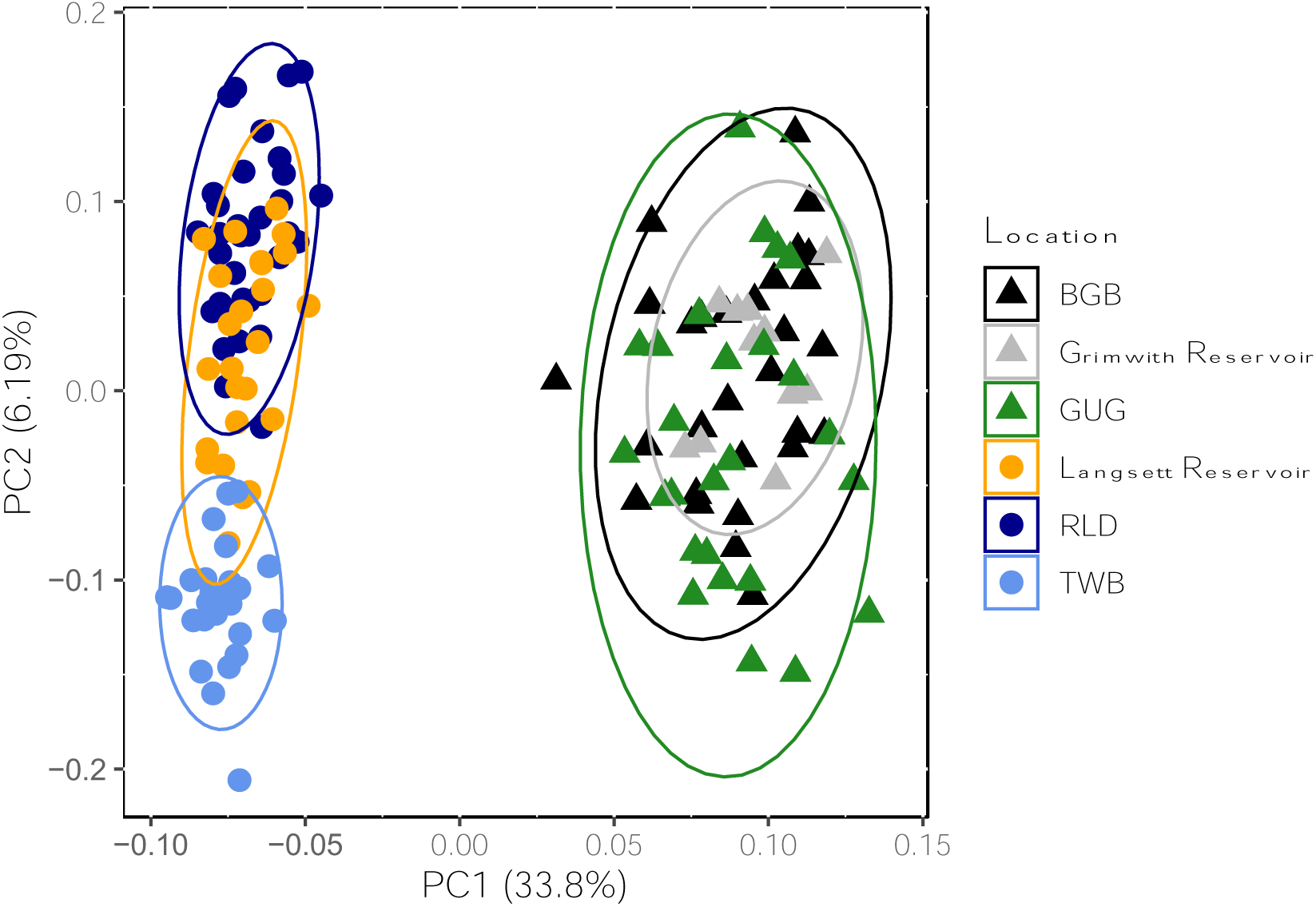
Genetic structuring of brown trout populations from the Langsett (circles) and Grimwith (triangles) systems before the fish pass construction. The principal components analysis (PCA) plot is based on 81 SNPs. Dots represent individual fish, and variance (%) explained by the first and second axes are shown.

**Table 1:** Number of samples per location and year before and after the fish pass opening on the RLD in late 2017.

| Location |  |  |  |  |  |  |
| --- | --- | --- | --- | --- | --- | --- |
|  | BGB | Grimwith Reservoir | GUG | Langsett Reservoir | RLD | TWB |
| 2015 | 12 | 0 | 15 | 2 | 21 | 10 |
| 2016 | 7 | 0 | 5 | 5 | 8 | 16 |
| 2017 | 11 | 12 | 9 | 23 | 7 | 8 |
| **Tot. before** | **30** | **12** | **29** | **30** | **36** | **34** |
| 2018 | - | - | - | 11 | - | - |
| 2019 | - | - | - | - | 1 | - |
| 2020 | 3 | 4 | - | 6 | 1 | - |
| 2021 | 9 | 12 | 12 | 10 | 1 | 1 |
| 2022 | - | - | - | 12 | 39 | 11 |
| **Tot. after** | **12** | **16** | **12** | **39** | **42** | **12** |

### Genetic analyses

#### DNA extraction, genotyping and quality control

DNA was extracted from dry scales and ventral fin samples employing a modified Mu-DNA: Tissue protocol (Sellers et al., 2018) using a solid phase reversible immobilization (SPRI) magnetic bead capture method (adapted from Roland & Reich, 2012) to isolate high molecular weight DNA. The DNA samples were sent to LGC Ltd, UK, for genotyping. A custom 192 SNP array designed for brown trout (see Saint-Pé et al., 2019, for details) was used for data generation. 174 SNPs were successfully genotyped. Eight SNPs positioned in mitochondrial DNA were excluded from the dataset, leaving 166 available loci for genomic investigation. We then performed quality control (QC) and filtering of SNPs data in PLINK version 1.9 and 2.0 (www.cog-genomics.org/plink/1.9/ and www.cog-genomics.org/plink/2.0/; Chang et al., 2015; Purcell et al., 2007). SNPs were filtered for missing genotypes (PLINK 1.9. command: *--geno 0.2*) and minor allele frequency (*--maf 0*.*01*). Individuals with high rate of missing SNPs (*--mind 0.2*) were discarded from analyses. Replicates and putative twin samples were also removed using PLINK 2.0 (*--king-cutoff 0.354*). These QC steps resulted in the retention of 81 SNPs when fish from Langsett and Grimwith systems were analysed together, with minor allele frequency filtering causing the loss of 85 SNPs. For consistency, the same 81 SNPs were retained for the temporal investigation of individual reservoirs.

#### Genetic variation of brown trout populations in Langsett and Grimwith reservoir systems <u>before</u> the fish pass construction

To investigate the genetic structuring of brown trout populations in Langsett and Grimwith reservoir systems prior to the fish pass opening, a principal component analysis (PCA) was performed in PLINK 1.9 using individuals from the RLD (*n* = 36), TWB (*n* = 34), Langsett Reservoir (*n* = 30), BGB (*n* = 30), GUG (*n* = 29) and Grimwith Reservoir (*n* = 12). Using the same samples, ADMIXTURE v.1.3. (Alexander, Novembre & Lange, 2009) was used to infer the most likely number of genetic clusters (K, testing from K=1 to K=6), determined based on the lowest cross-validation error. To investigate whether brown trout residing in Langsett reservoir were spawning and self-recruiting, a further ADMIXTURE analysis (testing from K=1 to K=3) was carried out using only fish from Langsett.

#### Genetic variation of brown trout populations in Langsett and Grimwith reservoir systems <u>after</u> fish pass construction

To assess temporal changes in genetic structuring in Langsett and Grimwith systems, the samples from the two reservoir systems were then analysed separately using the same 81 SNPs used for preliminary analyses (see section above). First, a discriminant analysis of principal components (DAPC) using the ‘adegenet’ v. 2.1.10 R package (Jombart, 2008; Jombart & Ahmed, 2011) and ADMIXTURE analyses were performed to describe population structure over time in the two reservoir systems. In each reservoir dataset, samples were divided into six groups consisting of the sampling location (RLD, Langsett Reservoir, TWB and BGB, Grimwith Reservoir, GUG) and period of sampling (‘Before’ and ‘After’ fish pass opening in the RLD). The DAPC method was selected to maximise genetic differences among groups (‘Before’ and ‘After’ locations). This approach enabled robust detection of the presence or absence of genetic variation among these groups. Two PC axes were retained as predictors of genetic differences among these groups based on the parsimonious *k*−1 criterion (as opposed to the proportional variance criterion; Thia, 2023), where k is the number of sampling locations (three in our case). The ‘hierfstat’ v. 0.5.11 package (Goudet, 2005) was used to calculate pairwise *F_ST_* (Weir & Cockerham, 1984) among groups (*genet.dist* function, method = *WC84*) and observed heterozygosity (*H_o_*), gene diversity (*H_s_*) and inbreeding coefficient (*F_is_*) for each group (*basic.stats* function). Loci deviating from Hardy-Weinberg equilibrium were computed using the function *hw.test* in the ‘pegas’ v. 1.2 package (Paradis, 2010). The number of monomorphic loci was also assessed as a proxy of genetic diversity in each group using the software NeEstimator v. 2.1 (Do et al., 2014). Finally, to estimate the gene flow among groups, relative directional migration rates were evaluated based on *Nm* distance (i.e. effective number of migrants; Alcala et al,. 2014) using the *divMigrate* function in the ‘diveRsity’ v. 1.9.90 package (Keenan et al., 2013; Sundqvist et al., 2016). The scripts used to run the analyses are publicly available at: https://tinyurl.com/trout-genetics.

## Results

### Genetic variation of brown trout populations in Langsett and Grimwith reservoir systems <u>before</u> the fish pass construction

Brown trout in the two reservoir systems were genetically differentiated from one another, as shown by the first Principal Component (PC1) (Fig. 2). In the Grimwith (control) system, fish from the reservoir and “no barrier” tributaries (BGB and GUG) overlapped widely showing no genetic structuring within the system. This was confirmed by ADMIXTURE analysis, which identified only one genetic unit for the Grimwith system (Suppl. Fig. 3), and by the low pairwise genetic distances (*F_ST_* between −0.0041 and 0.0037; Table 2) and high gene exchange (mean *Nm* = 0.82; Table 3) between locations. In contrast, Langsett system (impact) showed genetic structuring, with the two artificially fragmented tributaries (TWB and RLD) genetically distinct from each other (Figs. 2, 3, Suppl. Fig. 3). In the main Langsett Reservoir, some individuals exhibited admixed genetic profiles from both tributaries (RLD and TWB), indicating interbreeding between brown trout originating from these two sources (Figs. 2, 3, Suppl. Fig. 3). However, there were also individuals showing high similarity with the tributaries, suggesting they originated directly from these headwater locations. (Figs. 2, 3, Suppl. Fig. 3). *F_ST_* comparisons (Table 4) also supported these findings showing higher values between RLD and TWB tributaries (*F_ST_* = 0.1289) than between these two tributaries and Langsett Reservoir (*F_ST_* = 0.0109 and 0.0736, for comparison with RLD and TWB, respectively; Table 4). Estimates of gene flow among groups were in line with the above results, showing an overall low gene exchange in the Langsett system (mean *Nm* = 0.24; Table 5), with nearly absence of gene flow between two tributaries (RLD and TWB) with weirs at their mouths (*Nm* = 0.07; Table 5). Overall, Grimwith reservoir system had a higher genetic diversity than Langsett reservoir system (Table 6), showing higher *H_o_* (mean = 0.34 vs. 0.20), *H_s_* (mean = 0.37 vs. 0.20) and only few monomorphic loci (mean ≃ 2 vs. 35).

**Table 2.** Pairwise genetic distance between locations in Grimwith system before and after the construction of the fish pass as inferred by *F_ST_*.

|  | BGB Before | BGB After | Grimwith Res. After | Grimwith Res. Before | GUG Before |
| --- | --- | --- | --- | --- | --- |
| BGB After | 0.0066 |  |  |  |  |
| Grimwith Res. After | 0.0065 | 0.0100 |  |  |  |
| Grimwith Res. Before | -0.0041 | -0.0039 | -0.0105 |  |  |
| GUG Before | 0.0037 | 0.0168 | 0.0121 | 0.0007 |  |
| GUG After | 0.0094 | 0.0114 | 0.0123 | 0.0013 | 0.0115 |

**Table 3.** Estimates of gene flow (from rows to columns) of locations in the Grimwith system before and after the fish pass construction in the RLD at Langsett system computed using *Nm*.

|  | Grimwith<br>Res. Before | Grimwith<br>Res. After | BGB<br>Before | BGB<br>After | GUG<br>Before | GUG<br>After |
| --- | --- | --- | --- | --- | --- | --- |
| Grimwith Res. Before |  | 0.81 | 0.99 | 0.45 | 0.69 | 0.47 |
| Grimwith Res. After | 0.69 |  | 0.78 | 0.42 | 0.58 | 0.40 |
| BGB Before | 0.72 | 0.67 |  | 0.49 | 1 | 0.47 |
| BGB After | 0.56 | 0.45 | 0.67 |  | 0.48 | 0.42 |
| GUG Before | 0.55 | 0.54 | 0.98 | 0.33 |  | 0.45 |
| GUG After | 0.46 | 0.41 | 0.61 | 0.34 | 0.50 |  |

**Table 4.** Pairwise genetic distance between locations in Langsett system before and after the construction of the fish pass as inferred by *F_ST_*.

|  | RLD<br>Before | RLD<br>After | Langsett Res.<br>After | Langsett Res.<br>Before | TWB<br>Before |
| --- | --- | --- | --- | --- | --- |
| RLD After | 0.0331 |  |  |  |  |
| Langsett Res. After | 0.0100 | 0.0398 |  |  |  |
| Langsett Res. Before | 0.0109 | 0.0326 | -0.0007 |  |  |
| TWB Before | 0.1289 | 0.1332 | 0.0798 | 0.0736 |  |
| TWB After | 0.1389 | 0.1423 | 0.0864 | 0.0871 | 0.0263 |

**Table 5.** Estimates of gene flow (from rows to columns) of locations in the Langsett system before and after the fish pass construction in the RLD computed using *Nm*.

|  | Langsett<br>Res. Before | Langsett<br>Res. After | RLD<br>Before | RLD<br>After | TWB<br>Before | TWB<br>After |
| --- | --- | --- | --- | --- | --- | --- |
| Langsett Res. Before |  | 1 | 0.49 | 0.25 | 0.11 | 0.09 |
| Langsett Res. After | 0.88 |  | 0.52 | 0.20 | 0.09 | 0.07 |
| RLD Before | 0.54 | 0.66 |  | 0.26 | 0.07 | 0.06 |
| RLD After | 0.29 | 0.29 | 0.31 |  | 0.07 | 0.06 |
| TWB Before | 0.15 | 0.15 | 0.07 | 0.07 |  | 0.24 |
| TWB After | 0.10 | 0.12 | 0.06 | 0.05 | 0.24 |  |

**Table 6.**
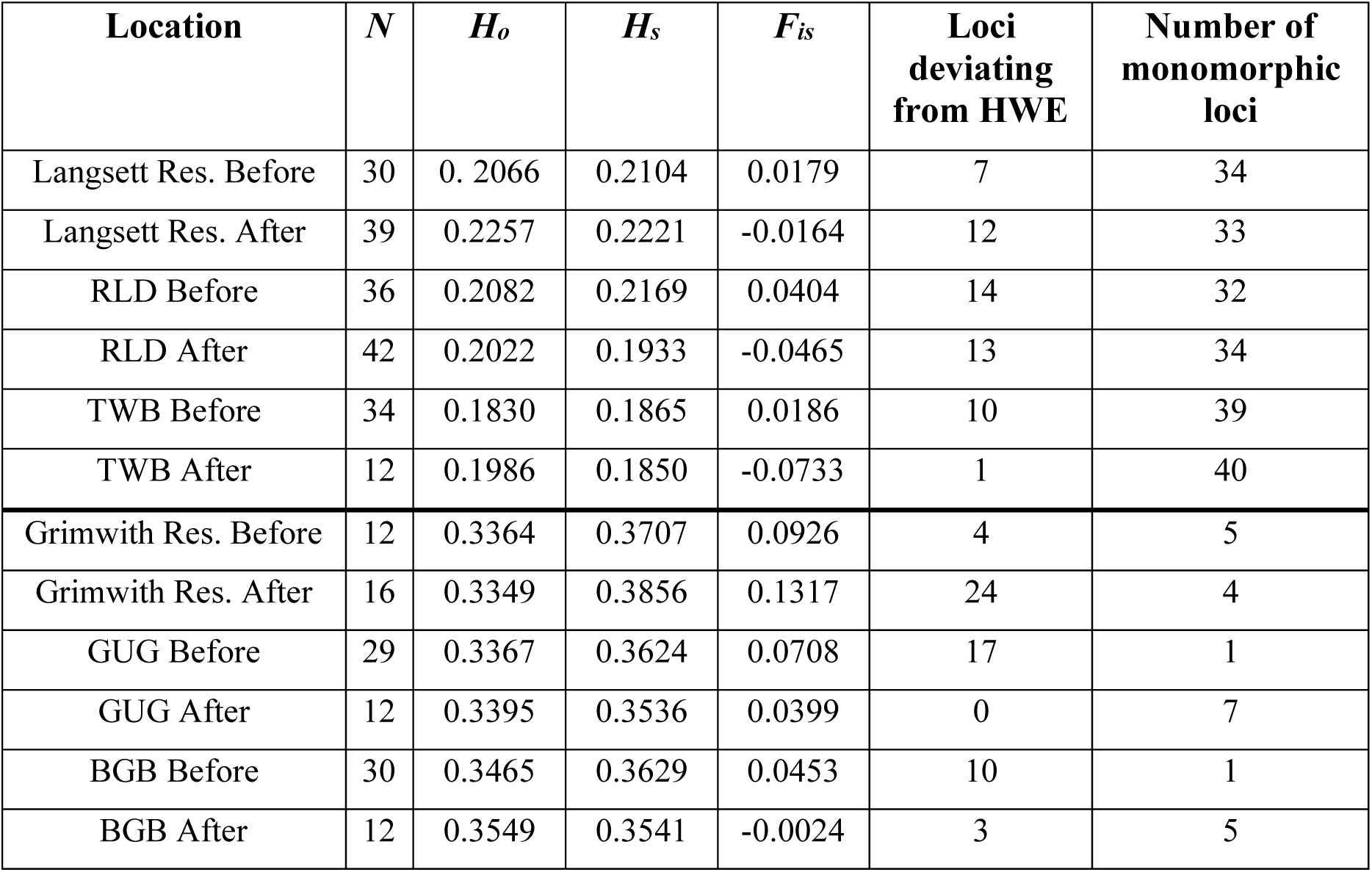
Estimates of observed heterozygosity (*H_o_*), observed gene diversity (*H_s_*), inbreeding (*F_is_*), loci deviating from Hardy-Weinberg Equilibrium (HWE) and number of monomorphic loci for each location in the Langsett and Grimwith systems before and after the construction of the fish pass in the RLD. Total number of loci = 81. *N* = number of samples.

| **Location** | **$N$** | **$H_o$** | **$H_s$** | **$F_{is}$** | **Loci deviating from HWE** | **Number of monomorphic loci** |
| --- | --- | --- | --- | --- | --- | --- |
| Langsett Res. Before | 30 | 0.2066 | 0.2104 | 0.0179 | 7 | 34 |
| Langsett Res. After | 39 | 0.2257 | 0.2221 | -0.0164 | 12 | 33 |
| RLD Before | 36 | 0.2082 | 0.2169 | 0.0404 | 14 | 32 |
| RLD After | 42 | 0.2022 | 0.1933 | -0.0465 | 13 | 34 |
| TWB Before | 34 | 0.1830 | 0.1865 | 0.0186 | 10 | 39 |
| TWB After | 12 | 0.1986 | 0.1850 | -0.0733 | 1 | 40 |
| Grimwith Res. Before | 12 | 0.3364 | 0.3707 | 0.0926 | 4 | 5 |
| Grimwith Res. After | 16 | 0.3349 | 0.3856 | 0.1317 | 24 | 4 |
| GUG Before | 29 | 0.3367 | 0.3624 | 0.0708 | 17 | 1 |
| GUG After | 12 | 0.3395 | 0.3536 | 0.0399 | 0 | 7 |
| BGB Before | 30 | 0.3465 | 0.3629 | 0.0453 | 10 | 1 |
| BGB After | 12 | 0.3549 | 0.3541 | -0.0024 | 3 | 5 |

### Genetic variation of brown trout populations in Langsett and Grimwith reservoir systems <u>after</u> fish pass construction

Genetic diversity (*H_o_*, *H_s_* and number of monorphic loci) in Langsett (impact) and Grimwith (control) reservoir systems did not change over time with respect to one another, with Grimwith showing higher genetic variation than Langsett both before and after the fish pass construction (Table 6). The DAPC showed contrasting patterns across reservoir systems (Fig. 4 A, B). In Grimwith, the ‘Before’ and ‘After’ samples overlapped widely meaning the genetic variation of tributaries and reservoir did not change over time. The mean proportion of successful reassignment of individual fish to their original sampling site was low (18.4 %; Suppl. Table 3) indicating a poor accuracy in distinguishing the different groups. The presence of a unique stable genetic unit in the Grimwith system over time was also supported by ADMIXTURE showing absence of structuring (K =1 most likely number of units) and the high level of relative migration rate (0.33 – 1, mean *Nm* = 0.58; Table 3) and low genetic distances (*F_ST_* max. = 0.0168; Table 2) among temporal groups. Conversely, in the Langsett system, the genetic composition of the RLD tributary changed after construction of the fish pass, as shown by the reduced overlap along the second discriminant function between fish from the ‘RLD Before’ and ‘RLD After’ groups (Fig. 4A), compared with fish from Langsett reservoir and TWB whose ‘Before’ and ‘After’ samples overlap extensively (Fig. 4A). This was reflected by the successful reassignment rate that was of 73.8 % for the ‘RLD After’ (Suppl. Table 3). Consistently, ADMIXTURE confirmed the increased structuring by identifying three genetic units, one of which was the ‘RLD After’ (Suppl. Fig. 4). Accordingly, estimated gene flow among sites was lower between fish from ‘RLD After’ and ‘RLD Before’, ‘Langsett Before’ and ‘Langsett After’ (*Nm* = 0.20 – 0.31) compared to gene flow among ‘RLD Before’, ‘Langsett Before’ and ‘Langsett After’ (*Nm* = 0.49 – 1; Table 5). *Nm* values between TWB and the other locations remained low over time (*Nm* max. = 0.11; Table 5). Finally, estimates of *F_ST_* were also consistent with a differentiation process occurring in the RLD, with *F_ST_* values between RLD and Langsett Reservoir shifting from *F_ST_* = 0.0109 ‘Before’, to *F_ST_* = 0.0398 ‘After’ the fish pass was built (Table 4). In contrast, there were no changes in genetic distance between TWB and RLD and Langsett Reservoir over time, indicating stability in gene flow among these sites in Langsett system (Table 4).

**Figure 3.**
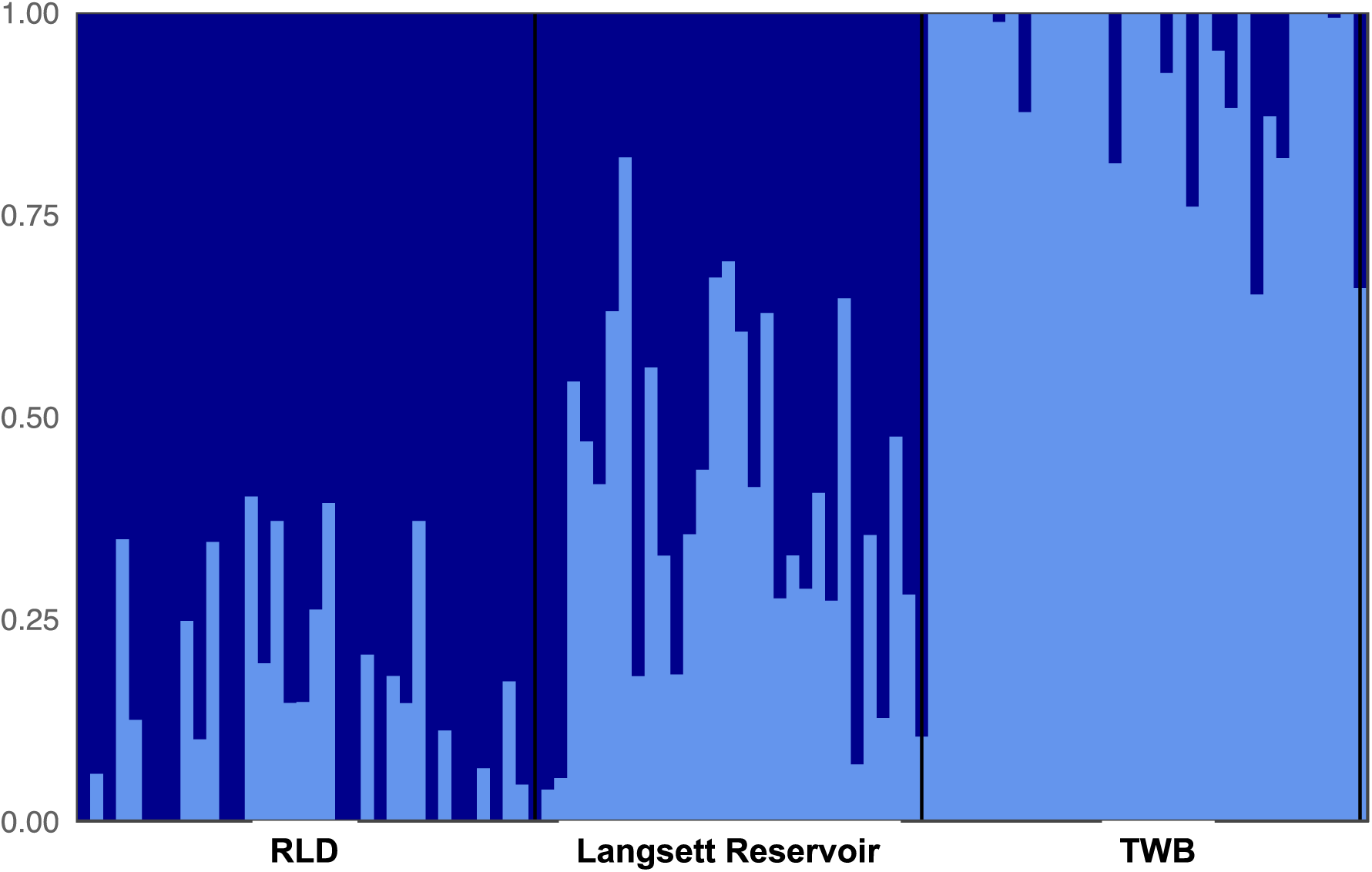
ADMIXTURE analysis plot (K=2) of brown trout populations from Langsett system before the fish pass construction. The analysis is based on 81 SNPs. Colours indicate different genetic clusters. Each fish individual is represented by a vertical bar.

**Figure 4.**
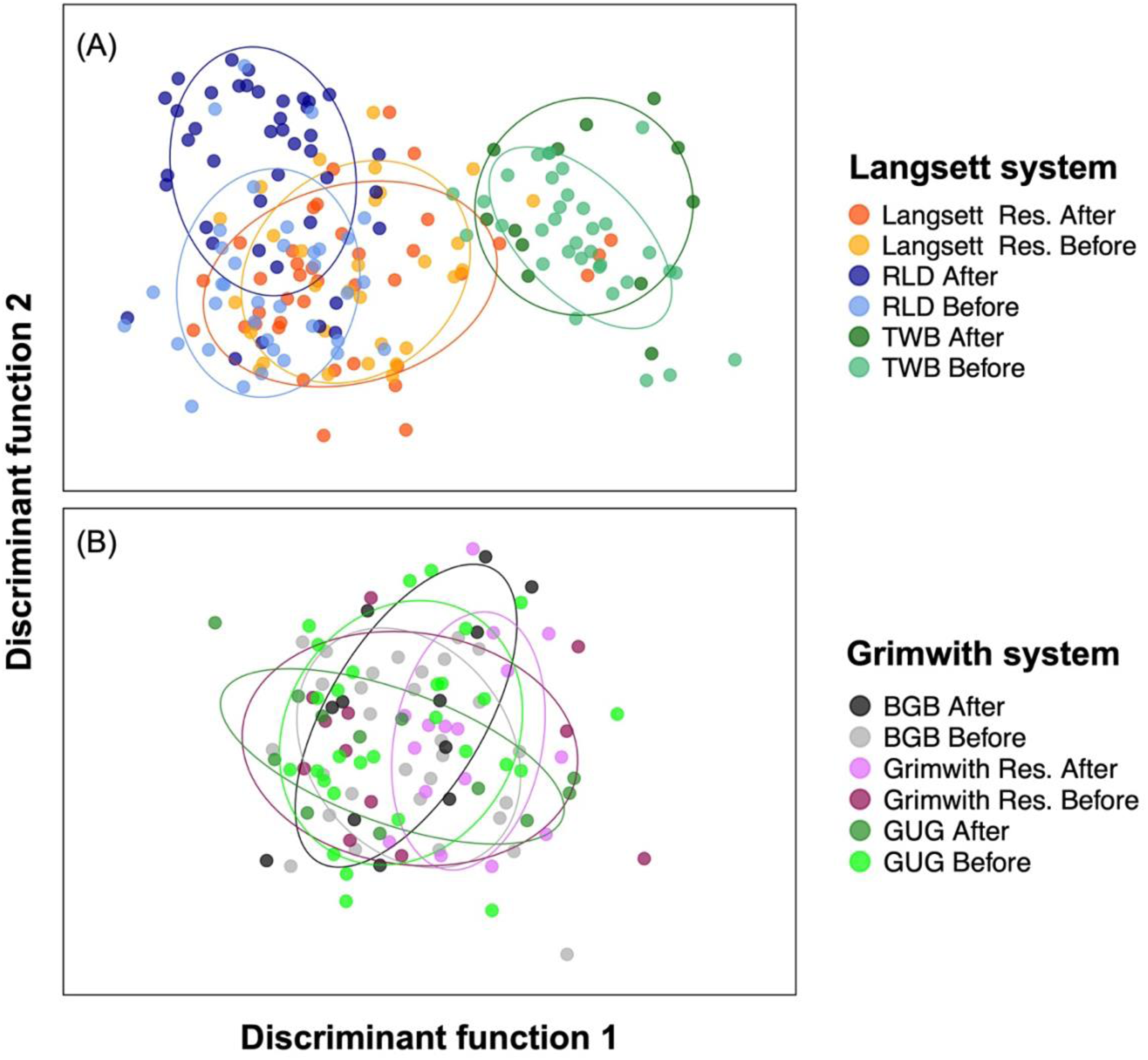
DAPC plot of Langsett (A) and Grimwith (B) systems before and after the fish pass construction in RLD. The plots are based on 81 SNPs. Dots represent individual fish.

## Discussion

Our study is one of the first to use before-after control-impact genetic samples for assessing the restoration of migratory corridors in freshwater systems and, to our knowledge, it is the first of its kind for brown trout.

Before the installation of the fish pass, we observed genetic structuring and reduced genetic diversity in the fragmented Langsett (impact) system compared to the “no-barrier” Grimwith (control) system. This is consistent with population genetics theory and previous studies on fish, where man-made river barriers reduced gene flow, which induced genetic erosion and structuring (Zarri et al., 2022). While genetic structuring *per se* does not necessarily imply negative consequences for populations, as it could indicate local fine-scale adaptation, which is common in salmonids (Garcia De Leaniz et al., 2007; Fraser et al., 2011), reduced genetic diversity poses risks for wild populations, potentially leading to maladaptation, decreased survival and fertility and increased vulnerability to environmental modifications (Hedrick & Miller, 1992; Lande, 1992; Hedrick & Kalinowski, 2000). It follows that brown trout in the Langsett system might be less resilient to future changes compared to those in the Grimwith system.

The genetic structure (PCA and ADMIXTURE) observed in the Langsett system prior to the fish pass construction indicates that brown trout were likely spawning within the reservoir. Indeed, some individuals caught in Langsett Reservoir showed admixture from the RLD and TWB tributaries, suggesting interbreeding in the reservoir but other individuals captured in the Langsett Reservoir seemed to be fish from the RLD and TWB tributaries, which is in accordance with telemetry analyses (Dodd et al., 2023). The higher genetic proximity between RLD and Langsett Reservoir (as opposed to TWB and Langsett Reservoir) is likely a result of the greater downstream migration into the reservoir from the RLD compared to the TWB (Dodd et al., 2023). The potential for spawning downstream of the RLD weir is supported by the presence of a short reach (50 m) of river with suitable substrate. They may also have spawned in the lentic water in the reservoir, as has been observed elsewhere for brown trout (Arostegui & Quinn, 2019). Conservation efforts could therefore focus on identifying and preserving possible spawning habitats in lakes and reservoirs in addition to mitigating riverine barriers. In this instance, fish pass construction was predicated on the assumption that brown trout must be provided access to lotic tributaries upstream of the lentic reservoir to complete their lifecycle and contribute to the next generation.

Despite the short period since the implementation of the fish pass on the RLD (∼ 5 years), changes in genetic structuring (but not genetic diversity) in the Langsett system were observed, confirming that restored connectivity can rapidly impact the genetic makeup of populations (Neville, Dauwalter & Peacock, 2016; Fraik et al., 2021; Prunier et al., 2023). Contrary to expectations though, genetic structuring increased, with the individuals in the RLD tributary becoming more differentiated from those in the Langsett Reservoir and TWB, rather than less. This genetic shift was supported by the DAPC, increased *F_ST_* and reduced inferred gene flow between RLD and Langsett Reservoir fish. The genetic profile of fish from TWB, which had no remediation measure, exhibited no changes. Previous studies on other salmonid species assessing genetic changes in response to river barriers removals, have shown contrasting patterns. Fraik et al. (2021) reported an increase in admixture, while Neville, Dauwalter & Peacock (2016) found no changes in genetic structuring, but an unexpected reduction in genetic diversity after barrier removal. Together with our findings, it seems that the impact of restored river connectivity on population genetics and migratory dynamics can be highly variable due to the complex interplay between evolutionary, ecological, and environmental processes. As a result, predicting these effects can be challenging, and thus further genetic analyses with BACI survey design are recommended to quantify the effects of river connectivity restoration efforts.

The differences in RLD after fish pass construction were attributed to juvenile fish in the most downstream survey site, suggesting they are the progeny of adult fish that have moved through the fish pass and spawned in the river (see Suppl. Figs. 1, 5). We propose that the fish pass has allowed fish that previously migrated downstream from the RLD into the reservoir the opportunity to return to their original home site upstream of the weir for spawning. This has reduced gene flow and promoted divergence between fish from the RLD and those from Langsett reservoir, as the latter are assumed to spawn and recruit downstream of the weir. Obviously, this hypothesis is reliant on the fact that brown trout home back to their natal place in the RLD for spawning, which has been suggested by telemetry analyses (Dodd et al., 2023). Previous studies have documented a wide range of homing accuracy in brown trout, ranging from fine-scale within-river homing to substantial straying among catchments (Halvorsen & Stabell, 1990; Armstrong & Herbert, 1997; Carlsson & Nilsson, 2000; Massa-Gallucci et al., 2010; Swatdipong et al., 2010; King et al., 2016; Mikheev et al., 2021; Källo et al., 2022, 2023). The potential for fine-scale homing behaviour and genetic admixing of brown trout above and below an impassable river barrier (a waterfall) has also been shown by Carlsson et al. (1999). *F_ST_* statistics revealed that fish captured downstream of the waterfall exhibited a closer genetic proximity to individuals sampled upstream of the barrier, compared to those captured at the end of a barrier-free adjacent tributary located also downstream of the waterfall. These findings support the notion of high philopatry among brown trout to the specific tributary and highlight the potential for genetic mixing with individuals descending from upstream regions in the presence of an impassable barrier as also found in other studies on salmonids (Meldgaard, Nielsen & Loeschcke, 2003; González-Ferreras et al., 2022).

An alternative explanation for the observed counterintuitive differentiation in the RLD tributary at Langsett could be genetic drift. This scenario may arise if the fish pass is ineffective, resulting in continued isolation of the RLD from the reservoir and subsequent directional genetic changes due to stochastic events. However, the absence of relevant genetic changes observed in both TWB (Langsett system) and the sites within the Grimwith system over the same time span, coupled with telemetry analyses of fish ascending the RLD fish pass (Dodd et al., 2023), makes this less likely and genetic diversity in the RLD (H_o_, H_s_ and proportion of monomorphic loci) remained constant over time. Alternatively, the genetic changes observed in the RLD could be attributed to fish migrating from the genetically distinct TWB population, traversing the reservoir, utilising the fish pass, and subsequently breeding in the RLD. However, if this were the case, we would expect to observe an increased genetic relatedness among these tributaries after the construction of the fish pass, which is not evident in our findings. Ideally, we would have also included a control reservoir that had tributaries both with and without barriers to migration to investigate genetic changes through time, but this was not possible. Kinship effects could have influenced the observed results, particularly considering the limited sample size in some groups, but this was mitigated by sampling across large reaches of river where possible.

Genetic differentiation driven by homing in salmonids is widely recognised for rapidly promoting the evolution of local adaptions (Hendry et al., 2000), even at microgeographic scale (Carlsson & Nilsson, 2000). Such local adaption can significantly increase the fitness of populations, highlighting the importance of managing wild fish populations for this (Fraser et al., 2011). Preserving genetically unique populations ensures the conservation of genetic and phenotypic traits that are well-suited to their specific habitats and contributes to the overall resilience and long-term viability of wild fish stocks. In this study, the availability of temporal (before-after) genetic samples allowed us to detect this incipient differentiation, which would not have been possible using other traditional methods used to assess the impact of fishways, such as telemetry and population abundance surveys. Since evolutionary processes are complex and can interact with a multitude of ecological and environmental factors, making observations over a longer time frame would help better understand the dynamics and outcomes of the ongoing genetic processes in this system.

Genetic divergence resulting from homing behaviour is an idea challenged by our findings in the no barrier reservoir system though. In Grimwith system, where migratory barriers are absent, complete genetic mixing, i.e. absence of homing behaviour, was observed over time. There are several potential explanations for this discrepancy between the two reservoir systems. These include variations in local ecological and environmental conditions, as well as differences in gene pools, which may trigger differential behavioural responses in the two systems. For instance, water chemistry differences between reservoir systems may trigger different homing behaviours, known to be driven by olfaction (Salmenkova, 2017). The origin of brown trout lineages and possible historical stocking in the two systems could also explain the observed dissimilarities, although there is no evidence to suggest stocking has occurred.

## Conclusions

This study demonstrates the efficacy of genetic tools in monitoring habitat fragmentation and restored connectivity. We were able to confirm that the isolated reservoir population was spawning within the reservoir before the addition of the fishway. Within a short period of five years after fish pass construction, we detected signals of genetic differentiation potentially driven by philopatric behaviour, that may lead to local adaptation. To our knowledge, this is the first study detecting such genetic shift in fish populations following restoration of migratory corridors. Continued monitoring these populations in the coming years will provide a better understanding of long-term population dynamics following restored ecological connectivity. Given the scarcity of similar studies, and the contrasting findings they present, we advocate for an increased use of genetic studies to monitor and manage wild populations, using a BACI design where possible. This allows managers to more holistically address whether the objectives of potentially expensive connectivity interventions have ultimately been successful.

## Data Archiving Statement

Data for this study are available at: to be completed after manuscript is accepted for publication.

## Acknowledgements

This study was funded by Yorkshire Water Services Limited. PM was supported by the Leeds-York-Hull Natural Environment Research Council (NERC) Doctoral Training Partnership (DTP) Panorama under grant NE/S007458/1. We also thank the University of Hull staff and post-graduate students for support in the field.

