## Supplementary_material for "Genetic consequences of improved river connectivity in brown trout (*Salmo trutta*, L.)"

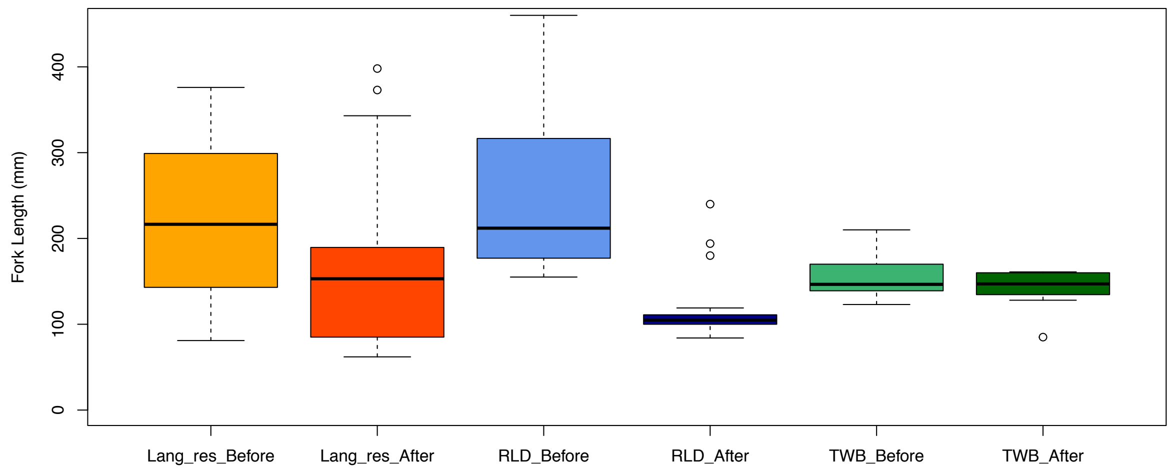


**Figure 1**: Size distribution of brown trout captured at Langsett reservoir system before and after the fish pass construction.

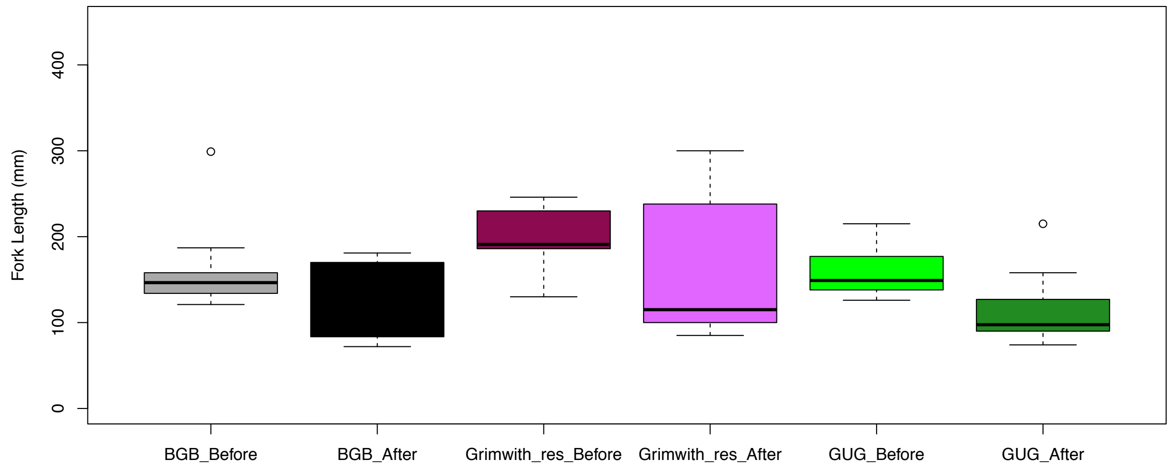


**Figure 2.** Size distribution of brown trout captured at Grimwith reservoir system before and after the fish pass construction.

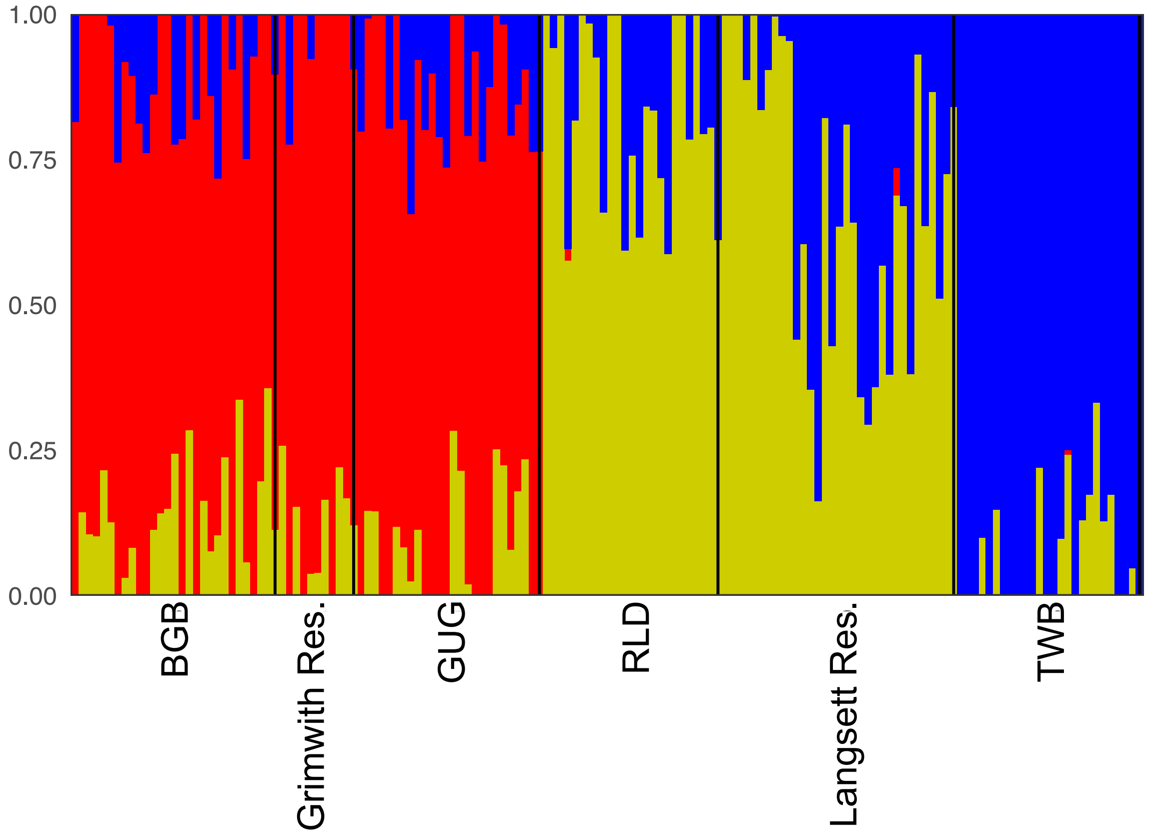


**Figure 3**: ADMIXTURE analysis plot (K=3) of brown trout populations from Langsett and Grimwith reservoir systems prior to fish pass opening on the RLD. The analysis is based on 81 SNPs. Colours indicate different genetic clusters. Each fish individual is represented by a vertical bar.

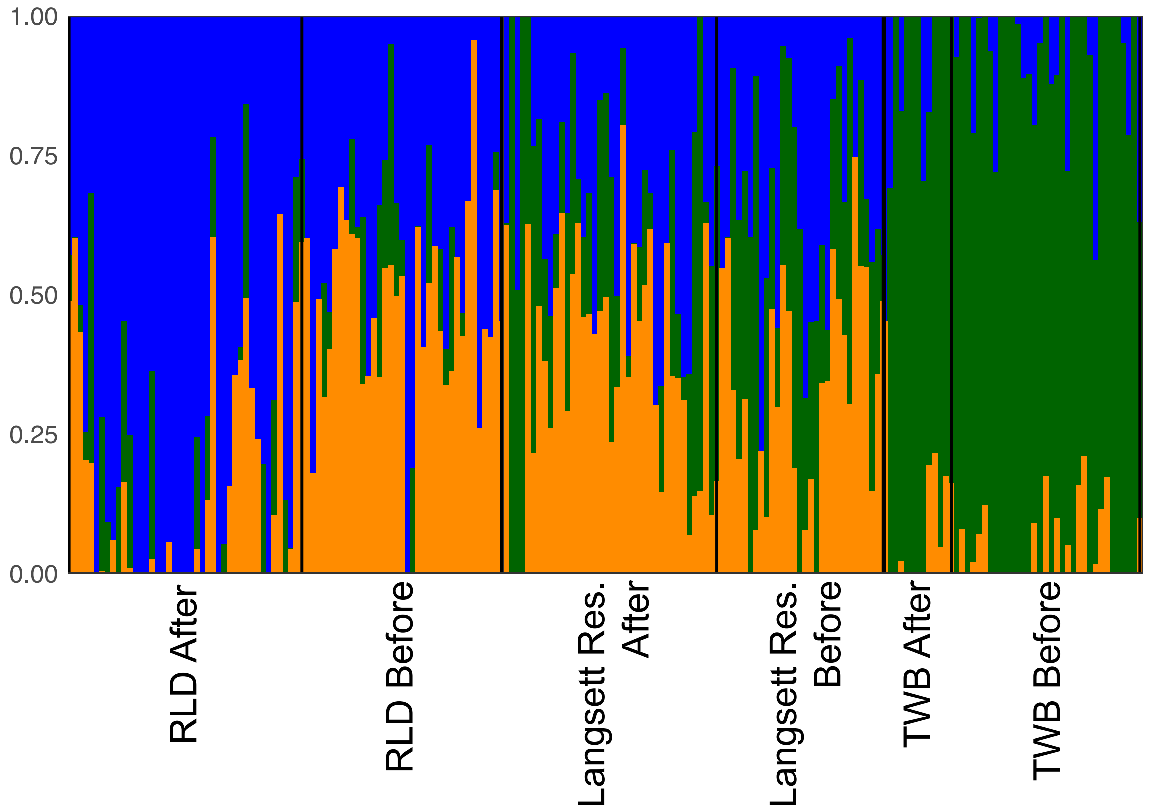


**Figure 4**: ADMIXTURE analysis plot (K=3) of brown trout populations from Langsett reservoir system before and after the fish pass opening on the RLD. The analysis is based on 81 SNPs. Colours indicate different genetic clusters. Each fish individual is represented by a vertical bar. An analogous analysis conducted on Grimwith reservoir system revealed absence of structuring within the system (K=1 most likely number of genetic units), therefore no figure was generated.

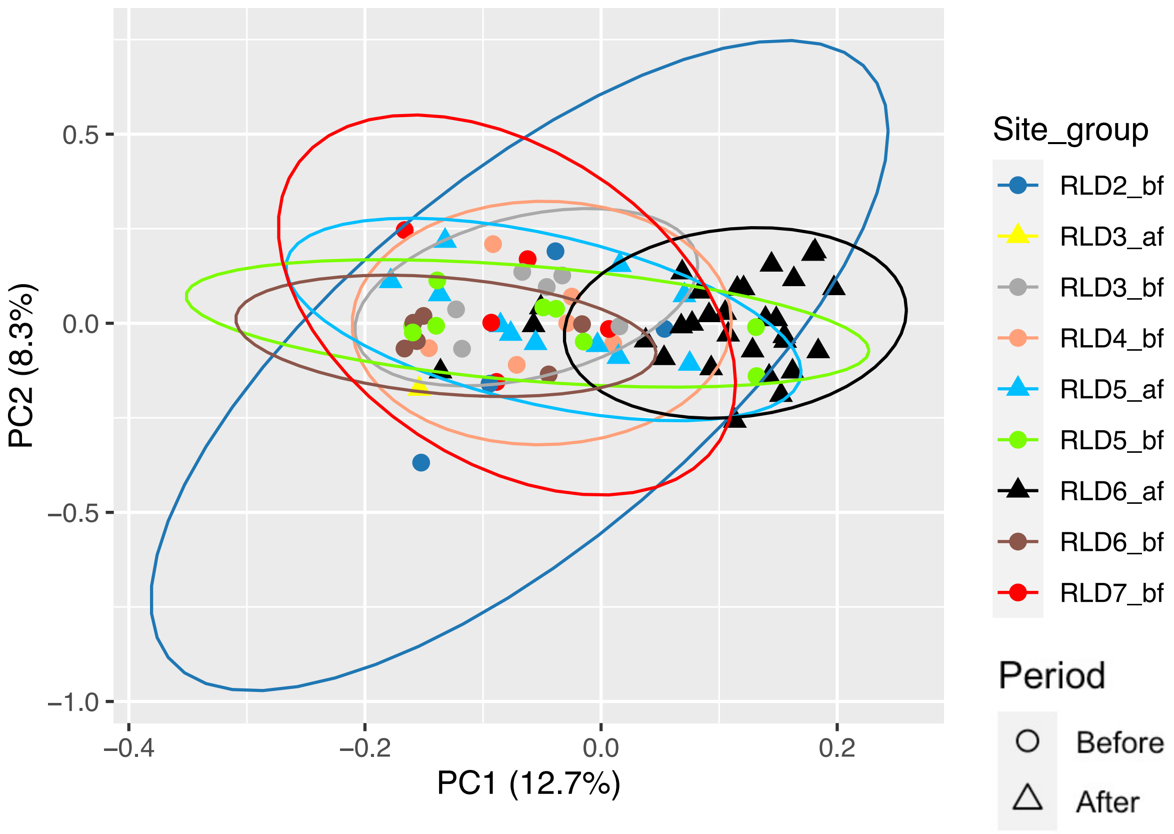


**Figure 5.** Genetic structuring of brown trout populations from the River Little Don (RLD) before and after fish construction. The principal components analysis (PCA) plot is based on 81 SNPs. Dots represent individual fish and are coloured based on sampling site and period (bf = before, af = after). Variance (%) explained by the first and second axes is shown.

**Table 1:** Samples caught at each sampling location at Langsett reservoir system with geographic coordinates.

| | | | Location | |
|---|---|---|---|---|
| **Lat.** | **Long.** | **Site ID** | **RLD Before** | **RLD After** |
| 53.49687 | -1.71360 | RLD2 | 4 | 0 |
| 53.49818 | -1.71359 | RLD3 | 6 | 1 |
| 53.49947 | -1.71274 | RLD4 | 6 | 0 |
| 53.50085 | -1.71101 | RLD5 | 9 | 11 |
| 53.50142 | -1.70834 | RLD6 | 6 | 30 |
| 53.50169 | -1.70611 | RLD7 | 5 | 0 |
| | | **Total** | 36 | 42 |
| | | | **Lang res Before** | **Lang res After** |
| Various sites | | Main reservoir | 13 | 6 |
| 53.50180 | -1.70163 | Downstream RLD weir | 16 | 32 |
| 53.49125 | -1.68776 | Downstream TWB weir | 1 | 1 |
| | | **Total** | 30 | 39 |
| | | | **TWB Before** | **TWB After** |
| 53.48475 | -1.68778 | TWB1 | 1 | 1 |
| 53.48586 | -1.68816 | TWB2 | 2 | 0 |
| 53.48730 | -1.68681 | TWB3 | 6 | 0 |
| 53.48825 | -1.68697 | TWB4 | 8 | 0 |
| 53.48925 | -1.68687 | TWB5 | 8 | 9 |
| 53.49007 | -1.68736 | TWB6 | 9 | 2 |
| | | **Total** | 34 | 12 |

**Table 2:** Samples caught at each sampling location at Grimwith reservoir system with geographic coordinates.

| | | | Location | |
|---|---|---|---|---|
| **Lat.** | **Long.** | **Site ID** | **BGB Before** | **BGB After** |
| 54.08865 | -1.92784 | BGB1 | 14 | 4 |
| 54.08816 | -1.92629 | BGB2 | 7 | 4 |
| 54.08724 | -1.92507 | BGB3 | 6 | 4 |
| | | NA | 3 | 0 |
| | | **Total** | 30 | 12 |
| | | | **Grim res Before** | **Grim res After** |
| Various sites | | Main reservoir | 12 | 16 |
| | | **Total** | 12 | 16 |
| | | | **GUG Before** | **GUG After** |
| 54.09082 | -1.91970 | GUG1 | 7 | 4 |
| 54.08961 | -1.91947 | GUG2 | 12 | 4 |
| 54.08822 | -1.91951 | GUG3 | 10 | 4 |
| | | **Total** | 29 | 12 |

**Table 3.** Proportions of successful reassignment (based on DAPC) of individual brown trout to their original groups.

| **Location** | **Proportion of successful reassignment** |
|---|---|
| BGB Before | 50.0 % |
| BGB After | 0 % |
| Grimwith Reservoir Before | 0 % |
| Grimwith Reservoir After | 18.8 % |
| GUG Before | 41.4 % |
| GUG After | 0 % |
| RLD Before | 52.8 % |
| RLD After | 73.8 % |
| Langsett Reservoir Before | 0 % |
| Langsett Reservoir After | 43.6% |
| TWB Before | 97.1 % |
| TWB After | 0 % |
